# Teleost fish can accurately estimate distance travelled

**DOI:** 10.1101/834341

**Authors:** C. Karlsson, J.K. Willis, M. Patel, T. Burt de Perera

## Abstract

Terrestrial animals compute shortcuts through their environment by integrating self-motion vectors containing distance and direction information. The sensory and neural mechanisms underlying this navigational feat have been extensively documented, but their evolutionary origins remain unexplored. Among extant vertebrates, the teleost fish make up one of the most diverse and earliest-branching phylogenetic groups, and provide a powerful system to study the origins of vertebrate spatial processing. However, how freely-swimming teleost fish collect and compute metric spatial information underwater are unknown. Using the Picasso triggerfish, *Rhinecanthus aculeatus*, we investigate the functional and mechanistic basis of distance estimation in teleost fish for the first time. We show that a fish can learn and remember distance travelled with remarkable accuracy. By analysing swimming trajectories, we form hypotheses about how distance is represented in the teleost brain, and propose that distance may be encoded by dedicated neural structures in a similar way to terrestrial vertebrates. Finally, we begin exploring the sensory mechanisms underlying distance estimation in fish. Many walking animals use a step counter for odometry. By quantifying finbeat use during our distance task, we show that a functionally equivalent finbeat counter is unlikely to provide reliable and precise distance information in an aquatic environment.

## 1 Background

A powerful way for an animal to navigate through its environment is through path integration. Self-movement vectors containing distance and direction information are constantly and automatically summated throughout any journey, providing the animal with an internal store of a vector taking it directly back to a starting position [1, 2]. The result is a dramatic increase in navigation efficiency - shortcuts can be constructed through entirely unexplored terrain, avoiding the need to use external information to retrace previous steps.

To achieve this navigational feat, an animal must have dedicated sensory mechanisms and neural structures to collect and process distance and direction information from self-motion cues. Such mechanisms have been extensively studied in terrestrial animals ranging from mammals such as humans [3] and rats [4, 5], and invertebrates such as spiders [6], ants [7], and bees [8, 9].

In contrast, how underwater species collect, process, and use metric spatial information to accurately navigate through their environment is largely unknown. An aquatic habitat is a fascinating environment to navigate through from a sensory and computational point of view. It contains sensory information not available on land, such as hydrodynamic cues from water flow, hydrostatic pressure from above, and electric currents carried through the water. Freely-swimming animals also have six degrees of freedom of movement (3 translational: forwards/backwards, left/right, up/down; 3 rotational: roll, pitch, yaw), compared to just three in surface-constrained animals (two translational: forwards/backwards, left/right; one rotational: yaw), which requires sensing and processing of spatial information in three-dimensions [10, 11, 12].

Among the animals facing these navigational challenges are the teleost fish, which display huge ecological diversity, inhabiting almost every aquatic niche on earth [13]. The teleost fish are therefore a valuable study system to shed light on how animals occupying aquatic environments have evolved to solve similar navigation problems to their terrestrial counterparts. As the most diverse and species rich vertebrate group, located in the sister clade to the tetrapods and lobe finned fish, studying how teleost fish encode metric spatial information is also important for understanding the representation of space in the vertebrate clade as a whole. There is increasing evidence to suggest that the teleost pallium is not only structurally equivalent, but homologous to the mammalian and avian hippocampus, placing the origin of a brain structure used in spatial memory as far back as 400 million years ago [14, 15, 16, 17]. However, in order to support this hypothesis, it must first be demonstrated at the behavioural level that the teleost fish possess similar navigation strategies to other vertebrates.

Early evidence from analysing swimming trajectories of fish trained to swim to a specific location to a gain a food reward suggests that fish can store an internal representation of distance and direction to a food reward relative to home [11], and compute such information in both horizontal and vertical space [12, 18]. The estimation of travel direction in the context of long range compasses has been studied in a range of migrating fish species, indicating some analogy with terrestrial and aerial species. Mosquitofish are able to orient using a time-compensated sun compass [19], while juvenile sockeye salmon and the Mozambique tilapia orient using magnetosensation [20, 21], and there is some evidence that rainbow trout use polarised light from above [22]. However, these abilities have never been directly quantified. Moreover, it is unknown whether teleost fish are able to measure distance travelled, which sensory mechanisms may support this behaviour, and the potential neural architecture that may underlie it. In this paper, we address these gaps by developing a behavioural paradigm to explore three key areas: (1) Can a teleost fish learn and remember distance information? (2) How might distance information be represented in the teleost brain? (3) What sensory cues do teleost fish use to collect distance information?

We develop a ‘match-to-sample’ behavioural task to assess whether fish can estimate travel distances. This is based on previous experiments done with the rat [5] and the desert ant [23]. The animal is trained to a given distance, and during testing we assess how accurately the animal can match this distance. We use the Picasso triggerfish (*Rhinecanthus aculeatus*) as our study species, typically found on shallow reef-flats throughout the Indo-Pacific Ocean. It has proven to be trainable in complex behavioural tasks and is naturally territorial so can be kept in isolated tanks in laboratory aquaria whilst maintaining natural behaviours [24, 25].

This behavioural paradigm can be used to form hypotheses about the mental representation, or neural encoding, of distance estimation in this fish species. In mammals, distance information is represented with remarkable accuracy by grid cells in the medial entorhinal cortex, receiving multiple sensory afferents from cortical brain regions [5, 26, 27, 28]. Whether teleost fish share this mechanism through common ancestry, or if they have evolved a separate architecture to represent distance information is unknown. It has been proposed that some animals may encode distance as a measure of travel time [8]. We test this latter hypothesis by exploring whether travel time is a good predictor of the variation seen across individual distance estimates. If travel time is used as a measure of distance travelled, the variance observed in the time and distance metrics would be equivalent, and we would observe faster swimming speeds for larger distance estimates. We demonstrate how this analysis, in combination with the distance estimation results observed, can be used to form hypotheses about the mental representation of distance in the teleost brain. These hypotheses can be tested into the future using neural lesioning studies and single cell recordings, whilst observing the behavioural output using the present behavioural task.

Finally, we explore how our study species acquires distance information using self-motion cues to compare how teleost fish have evolved to solve this problem in their aquatic world with animals walking on land. Previously studied invertebrates such as the desert ant, fiddler crab, and wandering spider partially or fully rely on an internal stride integrator as a measure of distance travelled through summation of inputs from leg mechanoreceptors [29, 30, 31, 32]. Humans are similarly able to estimate distance based on a function of walking speed, step length and step rate [33, 34]. The functionally equivalent mechanism for forward propulsion in a teleost fish would be the use of mechanosensory inputs from finbeat movements. The Picasso triggerfish uses three sets of fins for propulsion either in combination or in isolation: pectoral fins; undulating dorsal and anal fin pairs; and, the caudal fin. We investigate whether this species counts individual caudal finbeats (tailbeats), which provide the majority of rapid forward propulsion, or summates mechanosensory inputs across all finbeat combinations to measure distance travelled.

## 2 Methods

### 2.1 Subjects

Test subjects were five naive Picasso triggerfish, *Rhinecanthus aculeatus*, originating from coastal reefs on the Maldives, sourced through a local supplier. Individuals were housed in tanks measuring 0.45×0.30×0.75m (width x height x length) under a 12h/12h automated day/night flourescent light cycle, and provided with coral gravel, rocks and caves for enrichment. Salinity was kept constant at 35ppt using reverse osmosis water with added aquarium salts (Tropic Marine Centre Classic Sea Salt). Marine pellets (Ocean Nutrition Formula One Marine Pellet) and krill (Gamma Krill Pacifica) were provided as food rewards during training. Lance fish or cockles (Gamma) were fed as a supplement at the end of each training day. The experiment tank and home tanks were cleaned and water quality tested twice weekly. Ammonia and Nitrite were kept at 0ppm and Nitrate was maintained below 15ppm.

### 2.2 Behavioural Task

#### 2.2.1 Experimental set-up

A linear Perspex maze (supplementary fig. 1) measuring 0.25m high x 0.16m wide x 1.80m long (fig. 1) was built within a flow-through tank connected to the home water system to maintain constant water parameters. The walls and floor were patterned with regular black and white stripes of width 0.02m to provide basic visual contrast information, as many species have been shown to have impaired distance estimation abilities in the absence of optic flow [9, 23]. A perforated white screen was placed at either end to create laminar water flow whilst blocking the visual stimuli provided by cues external to the tank or the inlet and outlet pipes themselves. A moveable start area of dimensions 0.25m high x 0.16m wide x 0.30m long could be placed in one of three start area positions, all 0.1m apart (supplementary fig. 2). An infrared detector (SHARP 2Y0A21 proximity sensor) was then placed at water level overhead, 0.80m from the start area doorway. This was attached to an Arduino microprocessor, which through a Matlab (Mathworks Inc.) program controlled aquarium lights (Interpret Triple LED Lighting System, 0.75m) running along the top of the lateral maze walls. The voltage of the infrared (IR) detector varied between 0V and 3.5V depending on the strength of the reflection from objects which passed in front of it. We tested the response of the detector to objects in water and after this we set a threshold of 1.7V. As the fish passed beneath the detector, a voltage change was registered and when this exceeded 1.7V, the aquarium lights switched on. A Point Grey Grasshopper 3M camera (FLIR Machine Vision Cameras) was placed 1.1m above the water level to record testing trials.

**Figure 1:**
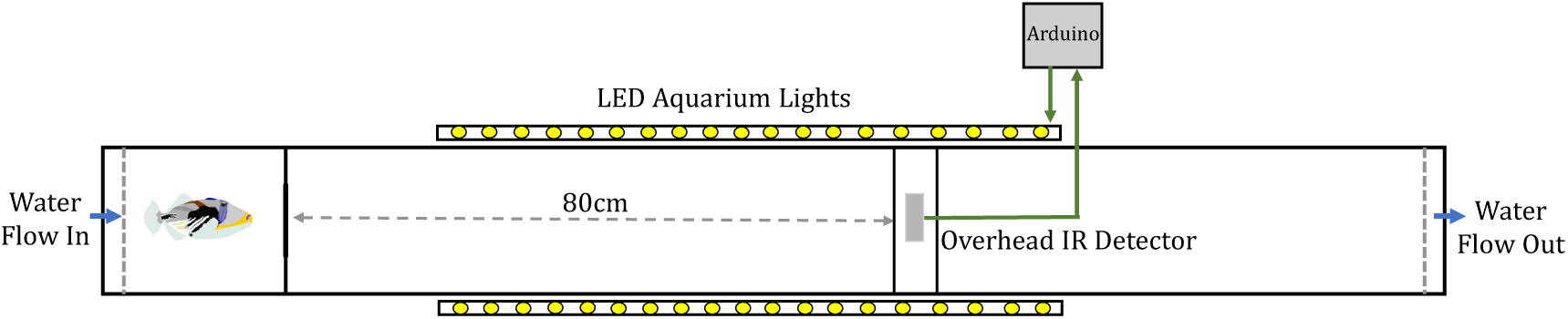
Distance estimation task basic training set-up. The linear maze was constructed inside a flow-through tank, with water flow in behind the start area (blue inlet arrow) and passive water flow out at the opposite end of the tank (blue outlet arrow). Perforated white screens separated the fish from the inlet and outlet pipes and ensured laminar flow. The maze walls and floor were lined with alternating black and white stripes of width 0.02m to provide constant optic flow information. The fish was placed in a movable start area and was trained to swim to an overhead infrared detector, which via an Arduino microprocessor caused the surrounding aquarium lights to switch on. The fish was trained to return to the start area for a food reward once the lights had switched on. An overhead camera recorded distance estimates during the testing phase.

#### 2.2.2 Distance Training

Fish were trained to swim 0.80m to the overhead infrared detector to switch on the aquarium lights and return home for a food reward. We trained the fish to pass beneath the detector to switch on aquarium lights to encourage them to learn the association between active swimming of a certain distance and a food reward rather than beaconing directly to a landmark. Training sessions lasted 10 minutes, or until 10 correct trials were complete. To control for use of external landmarks, each session the start area was moved randomly between three positions, located increasingly distally through the tank by 0.10m increments (supplementary fig. 2). The infrared detector was moved accordingly to maintain the correct distance of 0.80m. Training was considered complete when the fish swam directly out to the light flash and back on 80 percent of trials within the 10-minute session time limit, across three consecutive sessions.

#### 2.2.3 Testing: Can teleost fish estimate distance?

Test sessions were of variable length according to individual differences in motivation, with a pseudo-randomly alternating training + testing trial structure. During training trials (fig. 2A), the IR detector was placed 0.80m from the start area as before and the fish was rewarded in the start area if it was correct. During testing trials (fig. 2B), the infrared detector was moved to a decoy position 1.30m from the start area. Moving the detector distally tested whether the fish had learned to swim the correct distance, or if it had learned to beacon to the infrared detector landmark. Sessions always began with a testing trial set-up and trial order was randomised thereafter. 15 testing trials were completed at each start area position, resulting in a total of 45 distance estimates per fish. To control for use of external cues, each testing session the start area was once again moved between the three starting positions whilst keeping the distance to the infrared detector constant.

**Figure 2:**
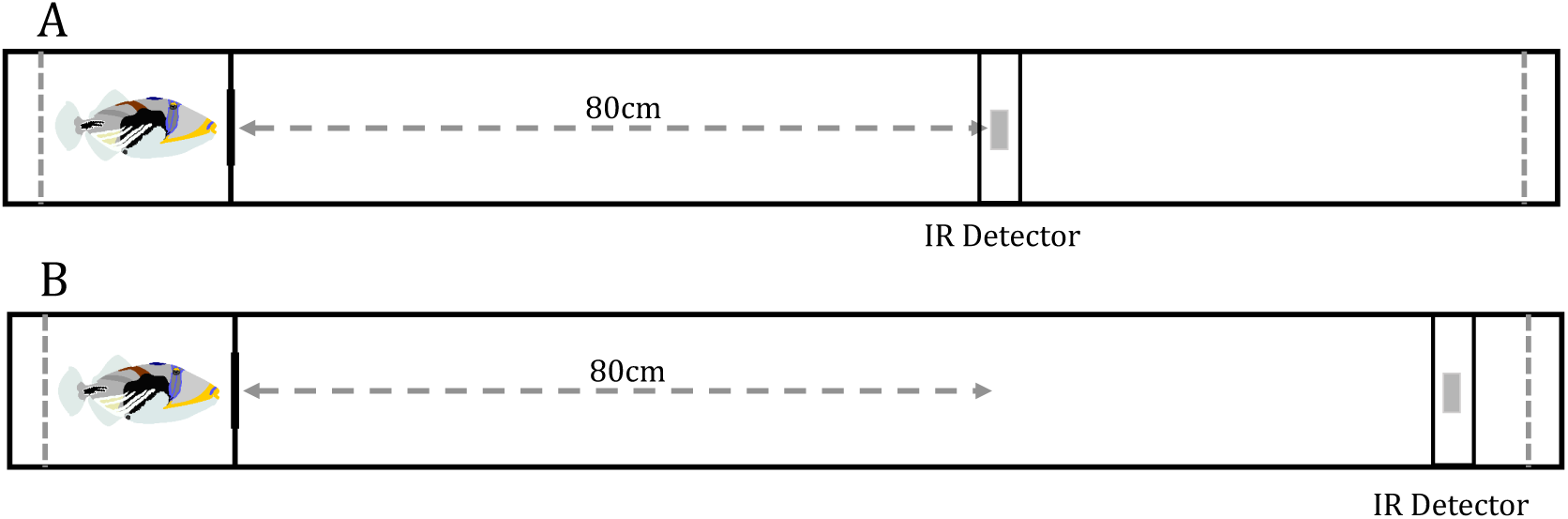
Distance estimation task training and testing set-up. (A) Training - the fish was trained to swim 0.80m to the infrared detector which when the fish passed beneath it, detected a voltage change and via the arduino computer caused the aquarium lights to switch on, signalling to the fish to return to the start area for a food reward. (B) Testing - during testing trials, the infrared detector was moved distally to 1.30m from the start area entrance. This was to test whether the fish had learned the correct distance, or to swim to the infrared detector landmark to encounter the light stimulus.

### 2.3 Analysis

Testing sessions were recorded using an overhead camera (Point Grey Grasshopper 3M) at 50 frames per second and saved as Audio Video Interleave (avi) files using the Streampix 7 video capture software (Image Width: 2448 pixels; Image Height: 350 pixels). Each testing trial was then extracted into a series of jpeg images using the Streampix 7 program making them compatible for analysis in a Matlab video tracking program.

#### 2.3.1 Extracting distance estimates

A distance estimate was considered the maximum point of the fish’s nose within a successful trial prior to turning home. A Matlab program was used to track the pixel coordinate position of the fish’s nose upon exiting the start area and the maximum point of the nose prior to turning home. Using the program R (The R Project, version 3.6.1) the total pixel distance travelled was calculated as the difference between the exit position and turning position, and estimates were converted to metric distances using the following conversion: 14.4 pixels = 0.01m.

#### 2.3.2 Extracting travel time

The frame number of the fish’s exit from the start area and the maximum point of the nose prior to turning home was recorded. The time taken was calculated as the difference between the exit and turning frame, and the frame rate of the video (50fps) was used to convert this to seconds per distance estimate.

#### 2.3.3 Finbeat analysis

Video recordings of the distance estimates were used to test whether the Picasso triggerfish uses proprioceptive inputs from finbeats to estimate distance. The Picasso triggerfish uses three sets of fins for propulsion either in combination or in isolation: pectoral fins; undulating dorsal and anal fin pairs; and, the caudal fin. The analysis was split into two levels. Caudal finbeats, or tailbeats, could be counted using the video data collected. Finbeats from the pectoral and dorsal/anal fin pairs could not be counted because they operate in such a way that makes a single beat difficult to distinguish. The total distance per testing trial dedicated to each finbeat type and combination was measured. Combinations included: caudal only; dorsal/anal pairs only; pectoral only; all finbeats; none (gliding); caudal and dorsal/anal; caudal and pectoral; dorsal/anal and pectoral. Information was first extracted as the total pixel distance for each finbeat type, and converted into a metric distance using the previous conversion (14.4 pixels = 0.01m).

## 3 Results

### 3.1 Teleost fish do have an internal representation of distance travelled

Figure 3 shows the average distance estimates for individual fish and the sample population (see also supplementary fig. 3 and table 1 for individual distance estimate distributions). All five fish avoided the overhead infrared detector in favour of turning at the perceived correct distance on all 45 testing trials. The average population level distance estimate was 0.803m (3sf) with a standard deviation of 0.0365m (3sf), which was not significantly different from the target distance of 0.80m - one-sample t-test, t_4,0.05_=0.0232 (3sf), p=0.983 (3sf).

**Table 1:** Comparing coefficients of variance for distance estimates and time taken. Coefficients were calculated for all five tested fish separately, calculated using the equation: (sample standard deviation/sample mean)x100. The ratio of coefficients was calculated using the equation: Time Taken Coefficient of Variance/Distance Estimate Coefficient of Variance.

| Fish | Distance Estimate<br>Coefficient of Variance | Time Taken Coefficient<br>of Variance | Ratio of Coefficients |
| --- | --- | --- | --- |
| A | 14.7 | 22.5 | 1.53 |
| B | 11.8 | 35.7 | 3.01 |
| C | 19.2 | 35.6 | 1.86 |
| D | 8.23 | 15.3 | 1.85 |
| E | 13.7 | 24.3 | 1.77 |

**Figure 3:**
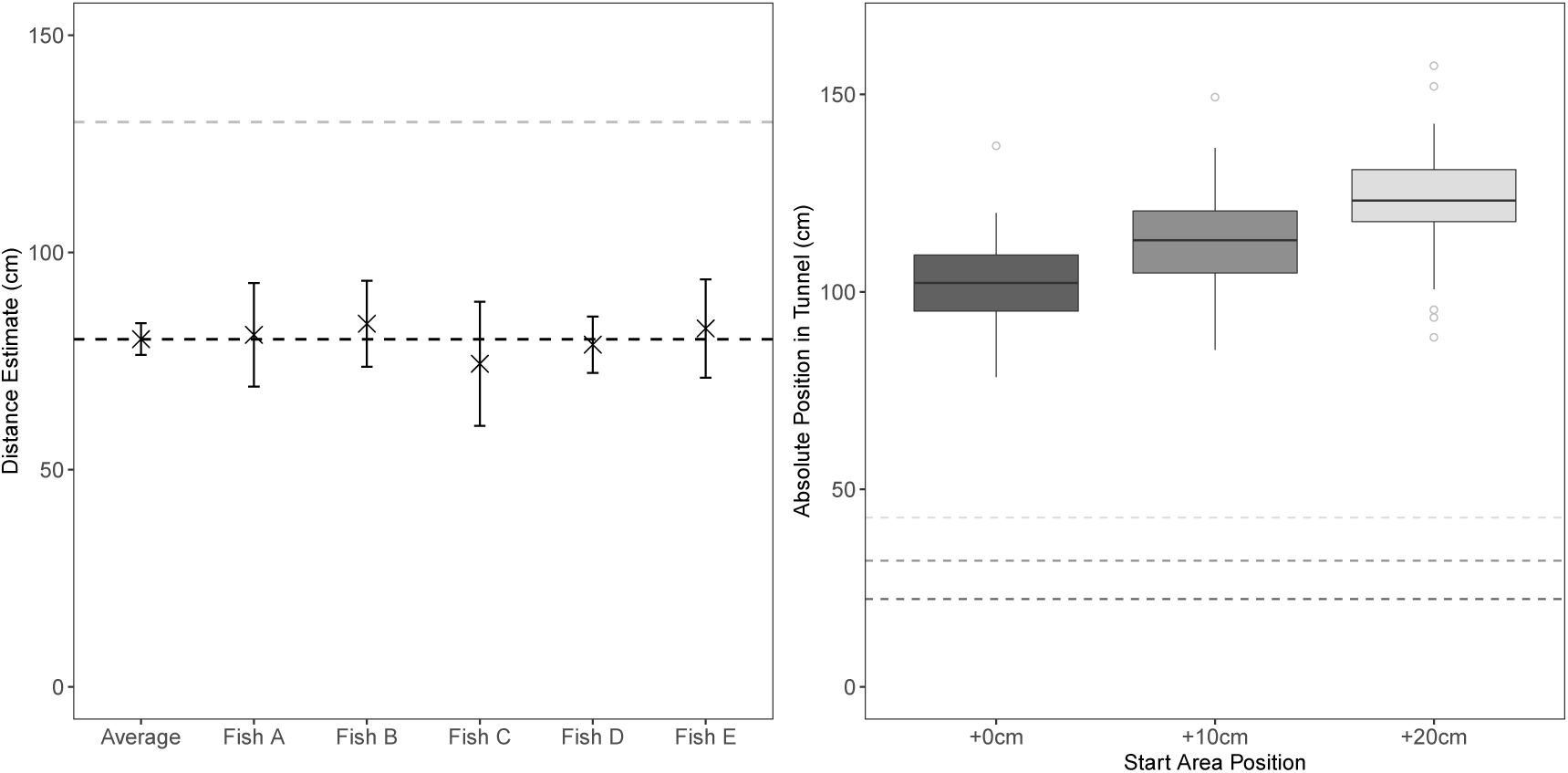
Left: Distance estimates produced by the population average and individual fish. Error bars indicate +/− 1 std dev. The black dashed line indicates the target distance (0.80m), and the grey dashed line indicates the decoy testing position of the overhead infrared detector controlling the lights. Right: Absolute distance estimate position within the tunnel maze for all fish, split by start area position. The start area moved between three positions: +0cm (dark grey), +10cm (mid-grey) and +20cm (light grey) - the dashed lines indicate the corresponding position of the start area doorway from the back of the tunnel.

Subjects were not using any additional positional cues internal or external to the maze itself to guide their turning points. Throughout training and testing, the start area was moved between three positions (identified as +0cm, +10cm, and +20cm) in order to shift the absolute position of the correct turning point in the tunnel while keeping the correct distance constant. If fish were generalising across positions and using a fixed external positional cue rather than an internal representation of distance, we would expect no difference in absolute turning point across the three start area positions. A linear mixed effects model (Absolute Estimate Point = Start Area Position (fixed effect) + Fish (random effect)) revealed start area position to be a good predictor of the absolute estimate point within the tunnel across fish (fig. 3, right): F_215,2_=69.8 (3sf), p*<*0.001, with absolute turning position for all three start area positions significantly different from each other (Tukey HSD pairwise comparison: 0:10 - t_215,0.05_=5.81, p*<*0.001; 10:20 - t_215,0.05_=6.00, p*<*0.001; 0:20 - t_215,0.05_=11.8, p*<*0.001). Individual fish identity explained some of the residual variation in distance estimates (Likelihood Ratio Test with and without random effect of fish = 7.63 (3sf), p=0.00575 (3sf), supplementary fig. 8). We therefore conclude that our fish are using an internal metric representation of distance travelled independently of external information.

### 3.2 Distance is not represented as a measure of travel time

To date, there is no evidence that any species uses travel time as a measure of distance. We tested this hypothesis for the distance estimates produced by our fish.

Travel time was a good predictor of distance travelled for our fish, with a positive relationship between the two at population level, fig. 4 - left (Linear Mixed Effects Model – Distance Estimate = Time Taken (fixed effect) + Fish Identity (random effect): t_219,0.05_= 9.16, p*<*0.001). A significant degree of residual variance was explained by the individual fish (Likelihood Ratio Test (model with and without the random effect of fish) = 43.9, p*<*0.001).

**Figure 4:**
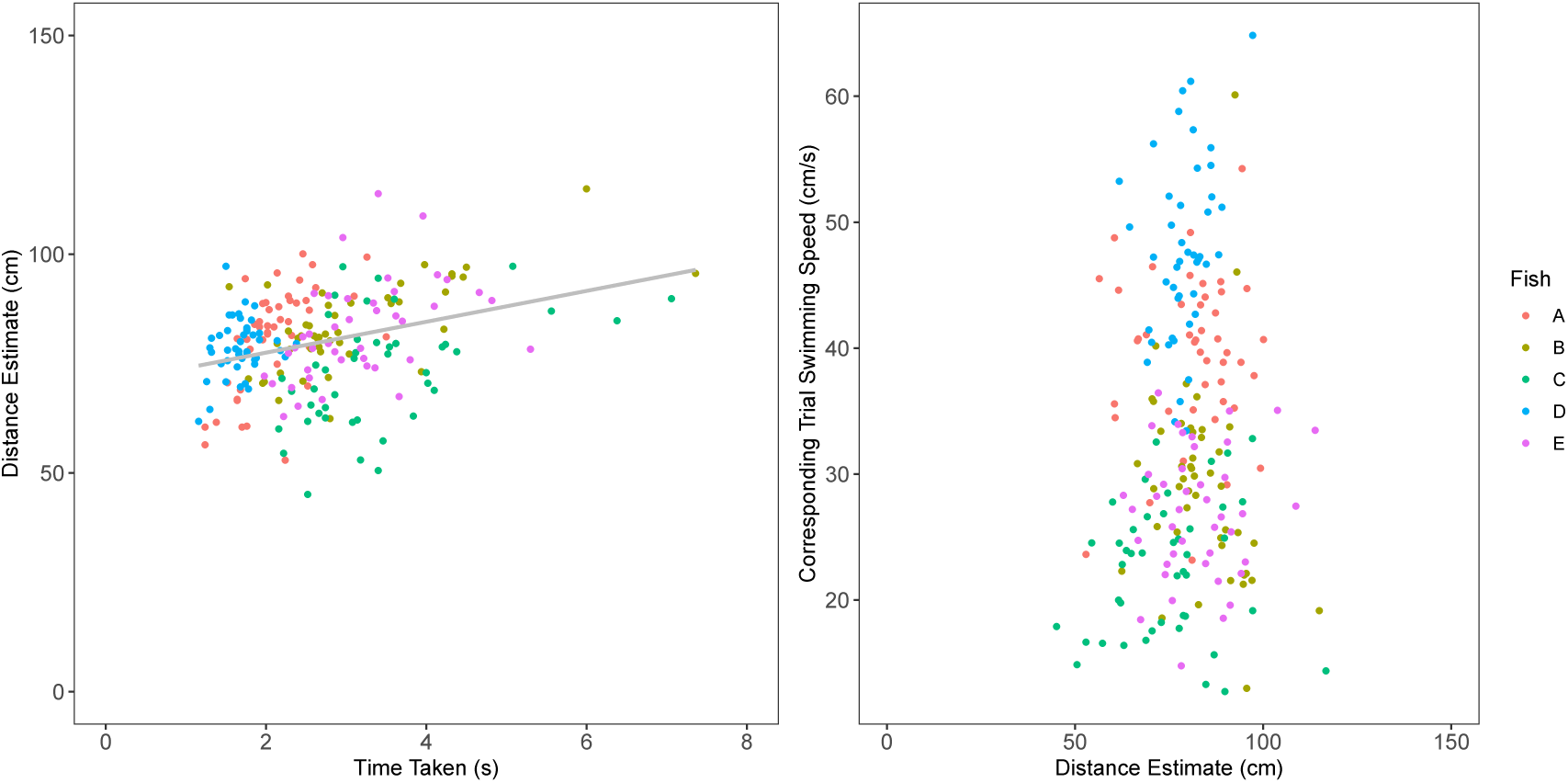
Left: Regression of Time taken against corresponding distance estimate. Equation of the linear regression line: Distance = 6.69 x Time taken + 61.9, t_219,0.05_= 9.16, p¡0.001. Right: Plot of distance estimate against corresponding trial swimming speed. There was no significant relationship between distance estimate and swimming speed: Distance = 0.0679 x Speed + 77.9, t_44,0.05_=0.694, p = 0.492. In both plots, coloured points represent individual fish identity.

This relationship is intuitive as if an individual is travelling at a near constant speed then we would expect to see larger distance estimates to take more time. The positive relationship would therefore emerge as a by-product of the error in their distance estimates. However, if time were to predict distance, then the coefficients of variation (measured as (standard deviation/average)x100) for each fish would be equal for both the time and the distance measurements. However, for all five fish the ratio of the coefficients was between 1.5 and 3, with a greater error for travel time than for the distance estimates produced (table 1).

We would also expect to see variation in swimming speed to be predicted by distance: if a fish was using time to measure distance, then further distance estimates would be associated with faster swim speeds. A linear mixed effects model was constructed as follows: Distance Estimate = Speed (fixed effect) + Fish (random effect). Speed was not a good predictor of distance estimate with no significant relationship between the two variables, fig. 4 - right (t_44,0.05_=0.492, p=0.492 (3sf)). Once more a significant degree of residual variance explained by the random effect of individual fish (Likelihood Ratio Test (model with and without the random effect of fish) = 9.06, p=0.00262(3sf)).

We conclude that Picasso triggerfish are therefore unlikely to represent metric distance as a measure of time travelled.

### 3.3 The Picasso triggerfish does not use finbeats to measure distance travelled

Video recordings of testing trials were used to explore whether the Picasso triggerfish uses mechanosensory inputs from finbeats as a sensory mechanism to collect information on distance travelled. Picasso triggerfish use caudal finbeats (here-on tailbeats) for rapid propulsion. They come in discreet units and can easily be counted from video recordings. A linear mixed effects model was constructed to test whether tailbeat number varied with distance estimates: Distance Estimate = Tailbeat Number (fixed effect) + Fish Identity (random effect). There was a positive relationship between tailbeat number and distance estimate, fig. 5, right (t_202,0.05_=5.52, p*<*0.001), with a significant degree of residual variation explained by the random effect of fish identity (Likelihood Ratio Test (model with and without the random effect of fish)=6.96, p=0.00835 (3sf)). However, there was a large variance in tailbeat number for each fish (fig. 5 – left, Range: Fish A, 3-14; Fish B, 3-10; Fish C, 0-14; Fish D, 5-14; Fish E, 3-14). Comparing the ratios of the coefficients of variance between distance estimates produced and the associated tailbeat number found the ratios varied from almost 2 (Fish A) to almost 4 (Fish C), table 2. The sensory information gathered from tailbeat number is therefore unlikely to provide the metric precision we see in the distance estimation data.

**Table 2:** Comparing coefficients of variance for distance estimates and tailbeat number. Coefficients were calculated for all five tested fish separately, calculated using the equation: (sample standard deviation/sample mean)x100. The ratio of coefficients was calculated usign the equation: Tailbeats Coefficient of Variance/Distance Estimate Coefficient of Variance.

| Fish | Distance Estimate<br>Coefficient of Variance | Tailbeats Coefficient<br>of Variance | Ratio of Coefficients |
| --- | --- | --- | --- |
| A | 14.7 | 27.7 | 1.88 |
| B | 11.8 | 29.8 | 2.52 |
| C | 19.2 | 81.2 | 4.23 |
| D | 8.23 | 20.7 | 2.51 |
| E | 13.7 | 30.9 | 2.25 |

**Figure 5:**
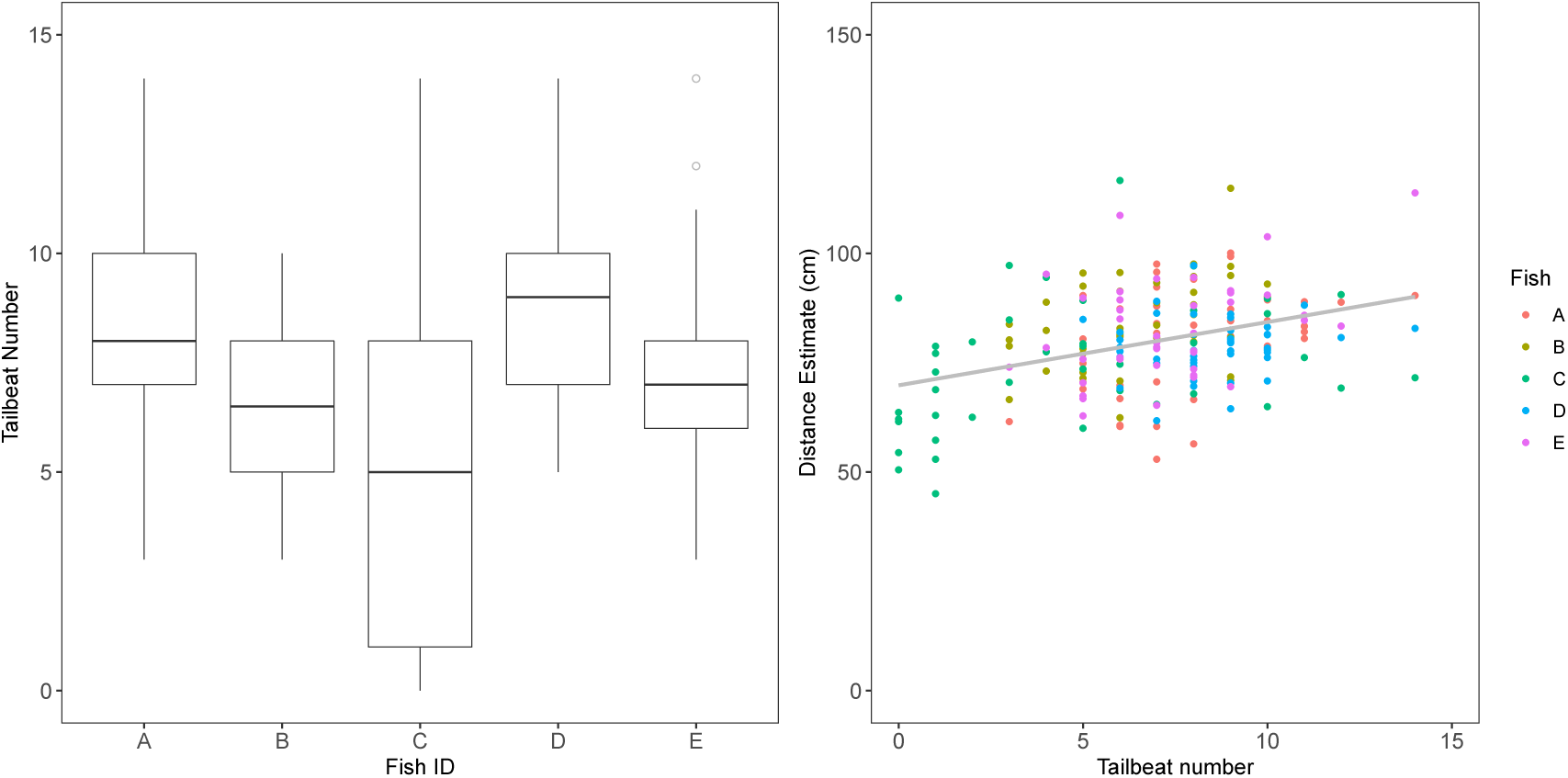
Left: Boxplot showing the range of tailbeat numbers for distance estimates for each fish (Fish A, 3-14; Fish B, 3-10; Fish C, 0-14; Fish D, 5-14; Fish E, 3-14). Right: Regression of tailbeat number against corresponding distance estimate for each fish (Distance Estimate = Tailbeats x 1.49 + 69.6, t_202,0.05_=5.52, p¡0.001). Coloured points indicate individual fish identity.

Tailbeat movements also only provide propulsion for a small fraction of the distance travelled for each estimate. The Picasso triggerfish exhibits three swimming modes using the caudal fin, pectoral fins and undulating dorsal and ventral fins together or in isolation for propulsion and steering. The distance travelled per swimming modality combination for each distance estimate was calculated. Combination categories were as follows: All fin types, caudal only, pectoral only, dorsal/ventral only, dorsal/ventral and pectoral, pectoral and caudal, dorsal/ventral and caudal, none (gliding). Results can be seen in fig. 6, revealing little consistency between trials both within and between individual fish. Tailbeats were often used in combination with pectoral fin movements and the undulating dorsal and anal fin pairs. There are also prolonged periods when the fish are not using tailbeats at all, instead relying on the other fin pairs for propulsion. Moreover, there are periods where the fish is gliding and no fins are moving. Even if the fish are able to sum the total mechanosensory inputs experienced across the fin types, this information is unlikely to provide the metric precision observed from the distance estimates.

**Figure 6:**
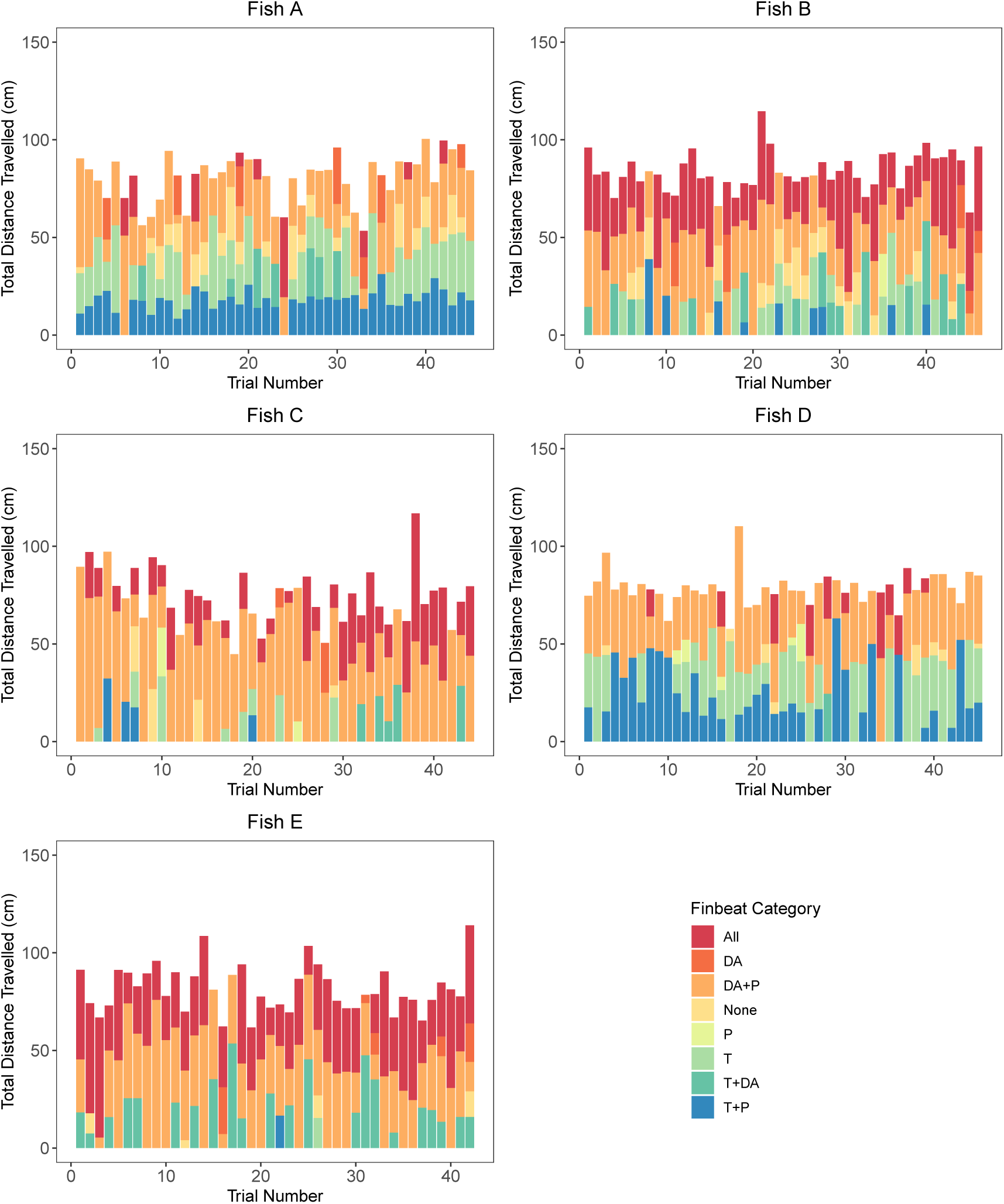
Bar plots showing the distance travelled using each finbeat type for each testing trial. Categories are as follows. All: All finbeat types used in combination, DA: Dorsal and Anal fins, DA+P: Dorsal, Anal and Pectoral fins, None: gliding - no fins used, P: Pectoral, T: Tail/caudal fin, T+DA: Caudal, Dorsal/Anal, T+P: Caudal + Pectoral.

## 4 Discussion

We show, for the first time, that teleost fish can estimate distance with remarkable accuracy (fig. 3). We explored how this distance information may be represented in the brain, and conclude that distance is not represented as a measure of travel time (fig. 4). Finally, we analysed video recordings of the distance estimates to begin investigating how Picasso triggerfish fish may acquire distance information. We tested whether the Picasso triggerfish uses mechanosensory inputs from finbeats to measure distance travelled, which would be functionally equivalent to using a step counter - a mechanism used by many vertebrate and invertebrate species walking on land. Based on the variance in finbeat types and number used, we conclude that the Picasso triggerfish is unlikely to use this mechanism to estimate distance (fig. 5, fig. 6).

Fish were trained to swim a distance of 0.80m to an overhead infrared detector. Passing beneath the detector switched on surrounding aquarium lights, which provided the cue to return to the start area for a food reward. During testing, this infrared detector was moved to a decoy position in the tunnel. The fish were therefore presented with a conflict: whether to favour the previously learned distance information and turn around at the previous position of the landmark cue, or to ignore the distance information and swim further to the new position of the landmark. Our fish consistently chose to ignore the novel position of the landmark cue in favour of returning to the start area once they had travelled the learned distance. We observed little deviation between individuals, producing a population average of 0.803m, (fig. 3 (left), supplementary fig. 3 and table 1). Each training and testing session, the start area moved systematically between three positions within the maze to control for use of absolute positional information. The fish were not using any absolute positional cues internal or external to the maze, as they did not generalise their turning point across these positions (fig. 3, right). We therefore conclude that the fish must be using a distance metric based on self-motion information. Ignoring the overhead landmark and not using any other external cues indicate that use of metric information acquired from self-motion may be prioritised by teleost fish when making navigation decisions. In an aquatic environment, landmarks may be scarce, hard to discern at a distance, or temporally unreliable. It may therefore be advantageous to prioritise use of internal metric information when making navigation decisions. Indeed, it has been demonstrated that reliance on external visual information declines for fish species and populations occupying environments with turbulent or variable flow [35].

We proceeded to explore how distance information could be encoded in the teleost brain. By extracting the time taken for each distance estimate from video recordings of test sessions, we tested the hypothesis that distance is encoded as a time metric. Although distance estimates did increase with time taken (fig. 4, left), the variability in travel time consistently surpassed the variability in distance estimates produced across all test subjects (table 1). Larger distance estimates were also not associated with faster swimming speeds, as would be expected if fish were using time as a measure of distance (fig. 4, right). This is consistent with results from previously studied vertebrate and invertebrate species. In the mammalian brain, spatial cells that are sensitive to elapsed travel time (time cells) have been discovered [36], but there is no evidence that such cell types underlie distance estimation. Indeed, humans tested in a similar way to our fish also exhibit greater error in time measurements compared to the distance estimates produced [34]. Among the invertebrates, honeybees also show no evidence of using travel time to estimate distance. Bees tested in a headwind underestimated distance travelled despite flights taking longer, and vice versa in a tailwind [37, 38]. For animals navigating in realistic, complex environments travel time is sensitive to disruption which reduces its reliability. For example, a Picasso triggerfish navigating through its coral reef habitat will have variable swimming speeds during any journey depending on whether it is in open sandy areas or enclosed reef areas, or whether it is alone or interacting with another animal. Time will therefore be progressing linearly, but distance travelled is likely to be non-linear and highly variable. The result is that travel time would not be a useful or reliable measure of distance.

Overall, our results indicate that a teleost’s mental representation of distance travelled is more accurate than a representation of travel time would support (table 1), if indeed travel time is represented in the teleost brain at all. The distance estimates produced by our fish are highly accurate, and must be supported by neural regions permitting precise representation of metric distance. In mammals, the most likely candidates for encoding distance estimation in mammals are the grid cells located in the medial entorhinal cortex. Rats with lesions to their MEC are unable to return to their home cage based on self-motion cues [4], and rats with medial entorhinal cortex lesions cannot estimate distance based on self-movement information [5]. Artificial agents trained to self-localise within a virtual environment using self-movement information spontaneously produced grid-like representations, convergent with the grid network found in mammals [39]. The authors argued that a grid cell like network provides the most parsimonious Euclidian map of space that enables vector based navigation. Our behavioural task produced similar results to an equivalent distance estimation task used to test distance estimation in rats [5]. However, whether a grid cell-like system also exists in the teleost fish, and if this would be a result of convergence or common ancestry is unknown. As neural manipulation and recording technologies have been developed for use in fish [40], our behavioural distance task now provides a valuable tool that can be used into the future alongside single cell recordings and lesioning studies to begin searching for brain regions and eventually cell types directly associated with distance estimation in teleost fish.

Finally, we explored the mechanosensory basis of distance estimation in our test species. We tested whether the Picasso triggerfish uses mechanosensory input from fin beats to measure distance travelled. Many terrestrial species use idiothetic information from a step counter for odometry. Humans are able to estimate distance travelled based on a function of walking speed, step length and step rate [33, 34]. Terrestrial invertebrates such as the desert ant, wandering spider and fiddler crab use proprioceptive inputs from slit sense organs on their legs internal stride integrators to measure distance ([29, 30, 31, 41]). We proposed that the functionally equivalent proprioceptive mechanism in fish would be the use of an internal finbeat counter. The Picasso triggerfish exhibits three swimming modes using the caudal fin, pectoral fins and undulating dorsal and anal fins together or in isolation for propulsion and steering. We primarily tested the role of tailbeat number in distance estimation, revealing a high variability in the number of tail beats across distance estimates (fig. 5). Comparing the coefficients of variance between tailbeat number and distance estimates show that tail beats alone are unlikely to provide the information needed to produce the precision seen in our distance estimate data (table 2). Trials with low tailbeat numbers are associated with increased use of the other two swimming modes, and during some trials fish use all three fin sets (fig. 6). There are also extended stretches during some, but not all, distance estimates where the fish glide through the water with no fin movements. We therefore suggest that even if the fish is capable of summing and integrating the total mechanosensory inputs from the different fin types, they are not using fin beat number to estimate distance. Unlike walking terrestrial species, where there is minimal ground movement as they walk across the surface, for fish swimming through variably moving water there is less likely to be a reliable directly proportional relationship between distance travelled relative to absolute space and the finbeat number used. This is especially true in rapidly moving bodies of water such as intertidal zones or fast-flowing rivers, where fin movements are often required to keep the fish stationary relative to the background [42]. The reliance on finbeats for odometry may therefore vary across fish species. Fish that occupy entirely motionless environments are not moving against or with currents may be more likely to experience a linear relationship between finbeat movements and distance travelled. Between-species variation in the sensory mechanisms used for odometry is observed among terrestrial animals. For example, while a step-counter is widely used by both vertebrate and invertebrate animals, flying honeybees which experience variable wind load instead rely almost fully on self-induced optic flow (the speed of visual motion across the retina) [9], and other species show varying reliance on energy use [43] and internal vestibular cues [5].

Future work should focus on unravelling species and habitat specific sensory mechanisms used for measuring distance travelled across the teleost clade. Different teleost species are likely to show a similar variation in sensory mechanisms to terrestrial animals, and this variation may be linked to their evolutionary ecology. As a coral reef fish occupying well-lit, spatially complex intertidal zones, the Picasso triggerfish may instead rely on visual information for odometry in a similar way to honeybees. In comparison, other less visual species such as blind cavefish may measure distance as a summation of proprioceptive inputs to their lateral line, and fish with an electric sense could measure distance using electrical currents in the water. Our behavioural task provides a robust paradigm through which we can continue to test these hypotheses.

## 5 Conclusion

Overall, we have demonstrated that teleost fish are able to estimate distance with comparable accuracy to terrestrial vertebrates, and they do so using self-motion cues alone. This distance information is not represented in the teleost brain as a measure of travel time, but we propose that it is more likely to be represented as a separate metric in the brain, perhaps in a similar way to mammalian grid cells. Our results indicate that it is unlikely that teleost fish use idiothetic information from finbeats to collect distance information, implying that the use of idiothetic cues from a stride integrator to estimate distance may have evolved in walking land animals alone. The behavioural task developed in this paper can be used into the future as a universal tool to explore the functional and mechanistic basis of distance estimation in fish at the neural and behavioural levels. Assessing distance estimation following neural lesioning and single cell recordings of candidate brain regions will allow us to study the neural mechanisms underlying distance estimation. By manipulating the sensory information provided during training and testing, we can also continue to explore the sensory mechanisms used in the odometers of different fish species. Gaining a more comprehensive understanding of how teleost fish encode metric space in their aquatic environment is a crucial next step in understanding the origin of metric spatial encoding in the vertebrate clade, and the navigational challenges faced by animals occupying an aquatic three-dimensional world.

## 6 Acknowledgements

We thank members of OxNav and in particular Tim Guilford for valuable input in the experimental design stages. This work was supported by the Biotechnology and Biological Sciences Research Council (BB/M011224/1) and a St John’s College Lamb and Flag studentship.

## References

[1] Etienne AS and Jeffery KJ. Path integration in mammals. Hippocampus, 14(2):180–92, Jan 2004. doi: 10.1002/hipo.10173.

[2] Etienne AS, Maurer R, Berlie J, Reverdin B, Rowe T, Georgakopoulos J, and Séguinot V. Navigation through vector addition. Nature, 396(6707):161–4, Nov 1998. ISSN 0028-0836. doi: 10.1038/24151.

[3] Loomis JM, Klatzky RL, Golledge RG, Cicinelli JG, Pellegrino JW, and Fry PA. Nonvisual navigation by blind and sighted: assessment of path integration ability. Journal of Experimental Psychology, 122(1):73–91, Mar 1993.

[4] Kim S, Sapiurka M, Clark RE, and Squire LR. Contrasting effects on path integration after hippocampal damage in humans and rats. Proceedings of the National Academy of Sciences of the United States of America, 110(12):4732–7, Mar 2013. doi: 10.1073/pnas.1300869110.

[5] Jacob P-Y, Gordillo-Salas M, Facchini J, Poucet B, Save E, and Sargolini F. Medial entorhinal cortex and medial septum contribute to self-motion-based linear distance estimation. Brain Structure and Function, 222(6):2727–2742, Aug 2017. doi: 10.1007/s00429-017-1368-4.

[6] Ortega-Escobar J and Ruiz MA. Visual odometry in the wolf spider Lycosa tarantula (Araneae: Lycosidae). The Journal of experimental biology, 217(Pt 3):395–401, 2014. doi: 10.1242/jeb.091868.

[7] Müller M and Wehner R. Path integration in desert ants, Cataglyphis fortis. Proceedings of the National Academy of Sciences of the United States of America, 85(14):5287–90, 1988.

[8] Harald W. Odometry and insect navigation. The Journal of experimental biology, 214(Pt 10):1629–41, May 2011. doi: 10.1242/jeb.038570.

[9] Srinivasan MV. Honeybees as a model for the study of visually guided flight, navigation, and biologically inspired robotics. Physiological reviews, 91(2):413–60, Apr 2011. doi: 10.1152/physrev.00005.2010.

[10] Jeffery KJ, Jovalekic A, Verriotis M, and Hayman R. Navigating in a three-dimensional world. The Behavioral and brain sciences, 36(5):523–43, Oct 2013. doi: 10.1017/S0140525X12002476.

[11] Holbrook RI and Burt de Perera T. Three-dimensional spatial cognition: freely swimming fish accurately learn and remember metric information in a volume. Animal Behaviour, 86 (5):1077–1083, 2013. doi: 10.1016/j.anbehav.2013.09.014.

[12] Burt de Perera T and Holbrook RI. Three-dimensional spatial representation in freely swimming fish. Cognitive processing, 13 Suppl 1:S107–11, Aug 2012. doi: 10.1007/s10339-012-0473-9.

[13] Near TJ, Eytan RI, Dornburg A, Kuhn KL, Moore JA, and Davis MP. et al. Resolution of ray-finned fish phylogeny and timing of diversification. Proceedings of the National Academy of Sciences of the United States of America, 109(34):13698–703, Aug 2012. doi: 10.1073/pnas.1206625109.

[14] Broglio C, Rodriguez F, and Salas C. Spatial cognition and its neural basis in teleost fishes. Fish and Fisheries, 4(3):247–255, Sep 2003. doi: 10.1046/j.1467-2979.2003.00128.x.

[15] Rodríguez F, López JC, Vargas JP, Broglio C, Gómez Y, and Salas C. Spatial memory and hippocampal pallium through vertebrate evolution: insights from reptiles and teleost fish. Brain Research bulletin, 57(3-4):499–503, 2002.

[16] Vargas JP, Rodriguez F, Lopez JC, Arias JL, and Salas C. Spatial learning-induced increase in the argyrophilic nucleolar organizer region of dorsolateral telencephalic neurons in goldfish. Brain research, 865(1):77–84, may 2000.

[17] Salas C, Broglio C, Durán E, Gómez A, Ocaña FM, and Jiménez-Moya F. et al. Neuropsychology of Learning and Memory in Teleost Fish. Zebrafish, 3(2):157–171, Jun 2006. doi: 10.1089/zeb.2006.3.157.

[18] Holbrook RI and Burt de Perera T. Separate encoding of vertical and horizontal components of space during orientation in fish. Animal Behaviour, 78(2):241–245, Aug 2009. doi: 10.1016/j.anbehav.2009.03.021.

[19] Goodyear CP and Ferguson DE. Sun-compass orientation in the mosquitofish, Gambusia affinis. Animal Behaviour, 17(4):636–640, 1969. doi: 10.1016/S0003-3472(69)80005-9.

[20] Shcherbakov D, Winklhofer M, Petersen N, Steidle J, Hilbig R, and Blum M. Magnetosensation in zebrafish. Current Biology, 15(5):R161–R162, 2005. doi: 10.1016/j.cub.2005.02.039.

[21] Quinn TP and Brannon EL. The use of celestial and magnetic cues by orienting sockeye salmon smolts. Journal of Comparative Physiology A, 147(4):547–552, 1982. doi: 10.1007/BF00612020.

[22] Hawryshyn CW, Arnold MG, Bowering E, and Cole RL. Spatial orientation of rainbow trout to plane-polarized light: The ontogeny of E-vector discrimination and spectral sensitivity characteristics. Journal of Comparative Physiology A, 166(4), Feb 1990. doi: 10.1007/BF00192027.

[23] Ronacher B and Wehner R. Desert ants Cataglyphis fortis use self-induced optic flow to measure distances travelled. Journal of Comparative Physiology A, 177(1):21–27, Jul 1995. doi: 10.1007/BF00243395.

[24] Cheney KL, Newport C, McClure EC, and Marshall NJ. Colour vision and response bias in a coral reef fish. The Journal of experimental biology, 216(Pt 15):2967–73, Aug 2013. doi: 10.1242/jeb.087932.

[25] Newport C, Green NF, McClure EC, Osorio DC, Vorobyev M, and Marshall N. et al. Fish use colour to learn compound visual signals. Animal Behaviour, 125:93–100, Mar 2017. doi: 10.1016/j.anbehav.2017.01.003.

[26] McNaughton BL, Battaglia FP, Jensen O, Moser EI, and M-B Moser. Path integration and the neural basis of the ‘cognitive map’. Nature reviews. Neuroscience, 7(8):663–78, aug 2006. doi: 10.1038/nrn1932.

[27] Moser EI and Moser M-B. A metric for space. Hippocampus, 18(12):1142–56, 2008. doi: 10.1002/hipo.20483.

[28] Moser EI, Kropff E, and Moser M-B. Place cells, grid cells, and the brain’s spatial representation system. Annual review of neuroscience, 31:69–89, Jan 2008. doi: 10.1146/annurev.neuro.31.061307.090723.

[29] Wittlinger M, Wehner R, and Wolf H. The ant odometer: stepping on stilts and stumps. Science, 312(5782):1965–7, Jun 2006. doi: 10.1126/science.1126912.

[30] Wittlinger M, Wehner R, and Wolf H. The desert ant odometer: a stride integrator that accounts for stride length and walking speed. The Journal of experimental biology, 210(Pt 2):198–207, Jan 2007. doi: 10.1242/jeb.02657.

[31] Walls ML, Layne JE, Altevogt R, von Hagen H-O, Zeil J, and Cannicci S. et al. Direct evidence for distance measurement via flexible stride integration in the fiddler crab. Current Biology, 19(1):25–9, Jan 2009. doi: 10.1016/j.cub.2008.10.069.

[32] Seyfarth E-A and Barth FG. Compound slit sense organs on the spider leg: Mechanoreceptors involved in kinesthetic orientation. Journal of Comparative Physiology, 78(2):176–191, 1972. doi: 10.1007/BF00693611.

[33] M-L Mittelstaedt and Mittelstaedt H. Idiothetic navigation in humans: estimation of path length. Experimental Brain Research, 139(3):318–332, Aug 2001. doi: 10.1007/s002210100735.

[34] Durgin FH, Akagi M, Gallistel CR, and Haiken W. The precision of locomotor odometry in humans. Experimental brain research, 193(3):429–36, Mar 2009. doi: 10.1007/s00221-008-1640-1.

[35] Odling-Smee L and Braithwaite VA. The influence of habitat stability on landmark use during spatial learning in the three-spined stickleback. Animal Behaviour, 65(4):701–707, 2003. doi: 10.1006/anbe.2003.2082.

[36] Eichenbaum H. Time cells in the hippocampus: a new dimension for mapping memories. Nature Reviews Neuroscience, 15(11):732–744, Nov 2014. doi: 10.1038/nrn3827.

[37] Srinivasan MV, Zhang S, and Bidwell N. Visually mediated odometry in honeybees. Journal of Experimental Biology, 200(19), 1997.

[38] Srinivasan MV, Zhang S, Lehrer M, and Collett T. Honeybee navigation en route to the goal: visual flight control and odometry. The Journal of experimental biology, 199(Pt 1): 237–44, Jan 1996.

[39] Banino A, Barry C, Uria B, Blundell C, Lillicrap T, and Mirowski P. et al. Vector-based navigation using grid-like representations in artificial agents. Nature, 557(7705):429–433, May 2018. doi: 10.1038/s41586-018-0102-6.

[40] Vinepinsky E, Donchin O, and Segev R. Wireless electrophysiology of the brain of freely swimming goldfish. Journal of Neuroscience Methods, 278:76–86, Feb 2017. doi: 10.1016/j.jneumeth.2017.01.001.

[41] Seyfarth E-A, Hergenrider R, Ebbes H, and FG Barth. Idiothetic orientation of a wandering spider: Compensation of detours and estimates of goal distance. Behavioral Ecology and Sociobiology, 11(2):139–148, Oct 1982. doi: 10.1007/BF00300103.

[42] Liao JC. A review of fish swimming mechanics and behaviour in altered flows. Philosophical transactions of the Royal Society of London. Series B, Biological sciences, 362(1487):1973–93, Nov 2007. doi: 10.1098/rstb.2007.2082.

[43] Proffit D, Stefanucci J, Banton T, and Epstein W. The role of effort in perceiving distance. Psychological Science, 14(2):106–12, Mar 2003.

