## Supplementary Information for "Teleost fish can accurately estimate distance travelled"

### 1 Methods

#### 1.1 Basic Set-up

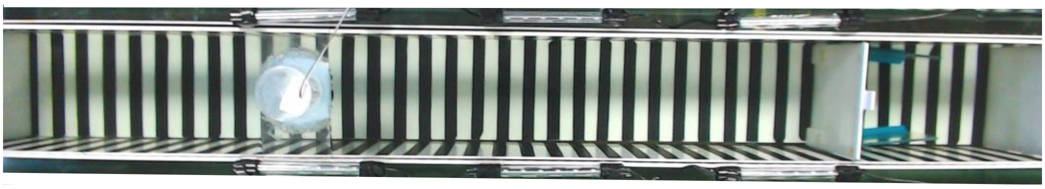

Figure 1: Overhead photo of main set-up in position +10. Two perforated white screens either end ensure laminar flow and prevent the fish from seeing outside of the tank. The start area is 30cm long, and the overhead infrared detector sits overhead 80cm from the start area doorway. This in turn controls aquarium lights running along the top of the maze walls symmetrically on either side.

Figure 1 is an overhead photo of the main set-up in position +20. Two perforated white screens ensure laminar flow and block visual cues external to the tank. The start area is 30cm in length and an overhead infrared (IR) detector is placed 0.80m from the start area doorway. The lateral and ventral black and white stripes provide basic optic flow information. Stripes are 0.02m wide and made from non-toxic waterproof black craft vinyl on white perspex. Two sets of aquarium lights run symmetrically along the top of the lateral maze walls (Interpret LED Lighting System 750mm, consisting of three 200mm light units arranged in series). When a fish passes beneath the IR detector, a voltage change above the threshold of 1.7V causes the arduino to turn on the aquarium lights until the fish turns around and returns home to receive a food reward.

#### 1.2 Controllling for use of external cues

The start area is moved between three different positions within the tunnel to prevent the fish from being able to use positional cues internal or external to the maze to learn the position of the infrared dececctor. Each session the start area and infrared detector move within the tunnel to a new position. These are as follows: +0 - the baseline positon with the rear perforated screen 0.02m from the rear of the tank; +10 - 10cm distal movement from the baseline +0 position; +20 - 20cm distal movement from the baseline +0 position. A schematic of these positions are found in fig. 2.

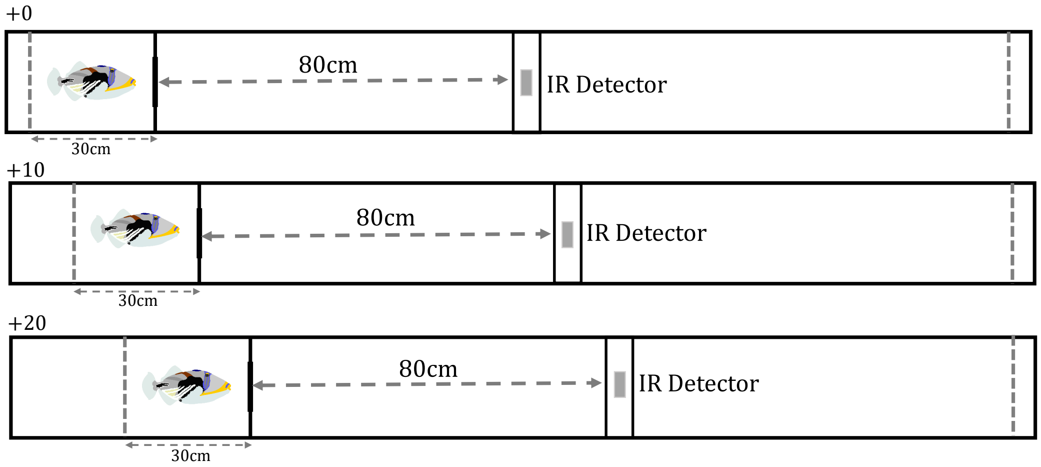


Figure 2: Schematic of the three start area positions. Each position is located increasingly distally through the tunnel by 0.10m increments (labelled +0, +10 and +20 to signify this). The rear perforated white screen moves accordingly to ensure the start area length is kept at 30cm. The infrared detector also moves accordingly to maintain the target distance of 0.80m.

| Fish | Average Distance Estimate (cm) | Standard Deviation |
|---|---|---|
| A | 81.0 | 11.9 |
| B | 83.5 | 9.90 |
| C | 74.4 | 14.2 |
| D | 78.6 | 6.50 |
| E | 82.6 | 10.8 |

Table 1: Average distance estimates and standard deviations for fish A-E (given to 3sf).

### 2 Individual fish distance estimates

Values of average distance estimates and standard deviations for all tested fish are in table 1, and individual distance estimate distributions are shown in fig. 3

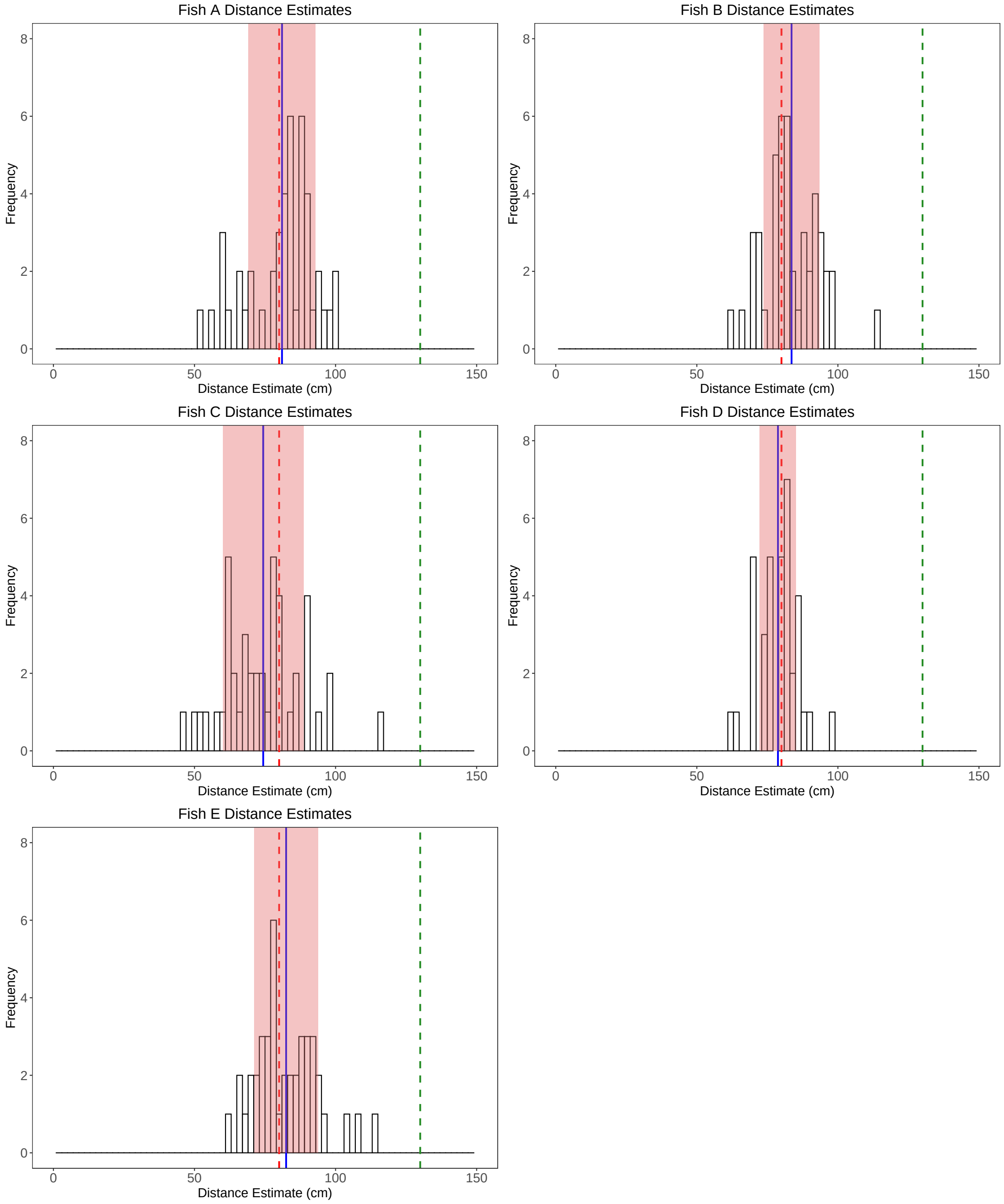


Figure 3: Distance estimate distributions for fish A-E. Average distance estimate = blue solid line, target distance (0.80m) = red dashed line, decoy position of infrared detector = green dashed line. Red shading indicates the standard deviations of the average distance estimate.

### 3 Testing assumptions

#### 3.1 t-test

Data points (average distance estimates, fish A-E), were independent and distance estimates were normally distributed (Shapiro-Wilk normality test, W = 0.925 (3sf), p=0.566 (3sf)), fig. 4. Parametric tests could therefore be used to test the distance estimates against the target distance of 80cm.

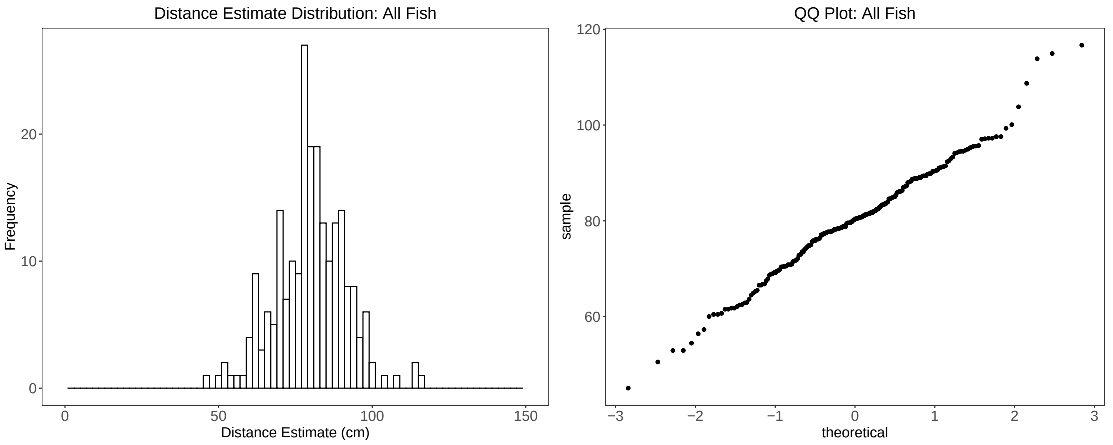


Figure 4: Normal QQ Plot providing an indication of the normality of distance estimates, allowing us to use parametric tests for analysis of distance estimates.

#### 3.2 Linear mixed effects model: Absolute Turning Positon = Start Area Position + Fish Identity

Testing the assumption of linearity is done by plotting the model residuals against the predictor (start area position). The resulting plot appears random (fig. 5, A), and we conclude the assumption has been met.

Variances were homogeneous (Levene's test, $F_{(212,2)} = 0.186$ (3sf), $p = 0.831$ (3sf)), supported by the even distribution of residuals plotted against the fitted values (fig. 5, B).

A QQ-Plot of the standardised residuals indicates no major deviation from the straight line, and we conclude that they are therefore normally distributed (fig. 5, C).

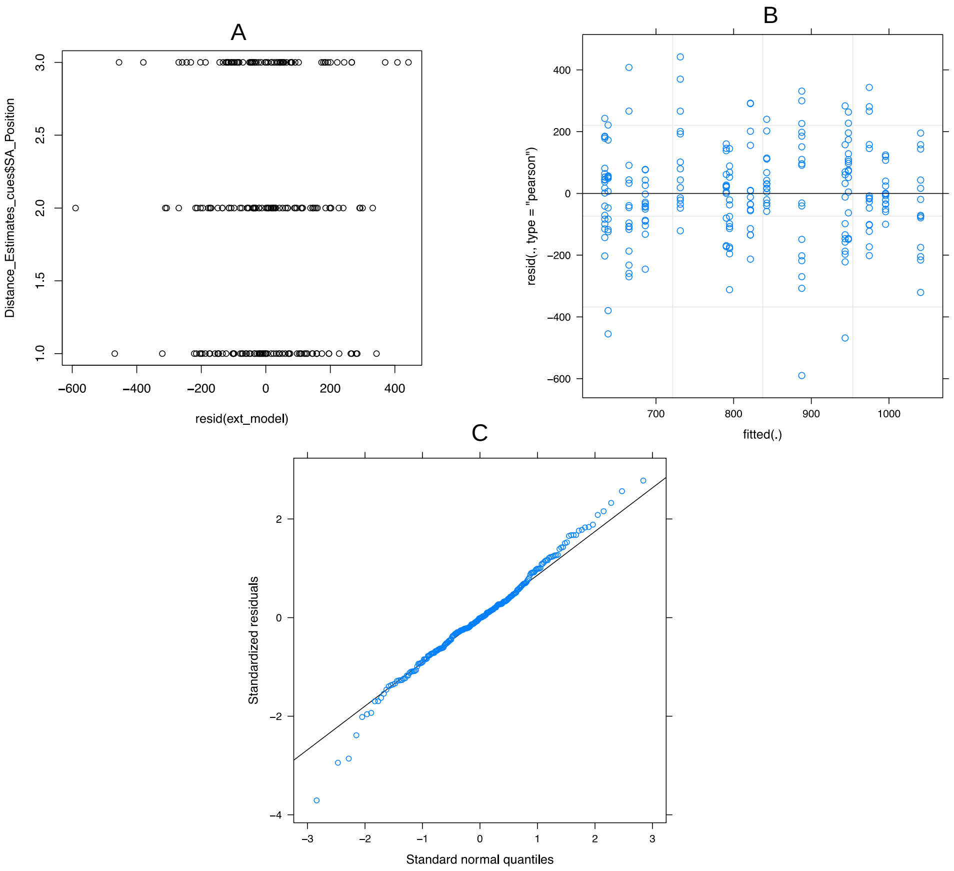


Figure 5: Testing mixed effects model assumptions (Absolute Turning Positon = Start Area Position + Fish Identity). (A): Plot of the model residuals against start area position. (B) A fitted vs residual plot revealing homogeneity of variance. (C)A QQ Plot of the standardised residuals, revealing a normal distribution.

##### 3.3 Linear mixed effects model: Distance Estimate = Time Taken + Fish Identity

The plot of residuals against the predictor (Time Taken) appears random (fig. 6, A), and we conclude the assumption has been met.

Variances were homogeneous, supported by the even distribution of residuals plotted against the fitted values (fig. 6, B).

A QQ-Plot of the standardised residuals indicates no major deviation from the straight line, and we conclude that they are therefore normally distributed (fig. 6, C).

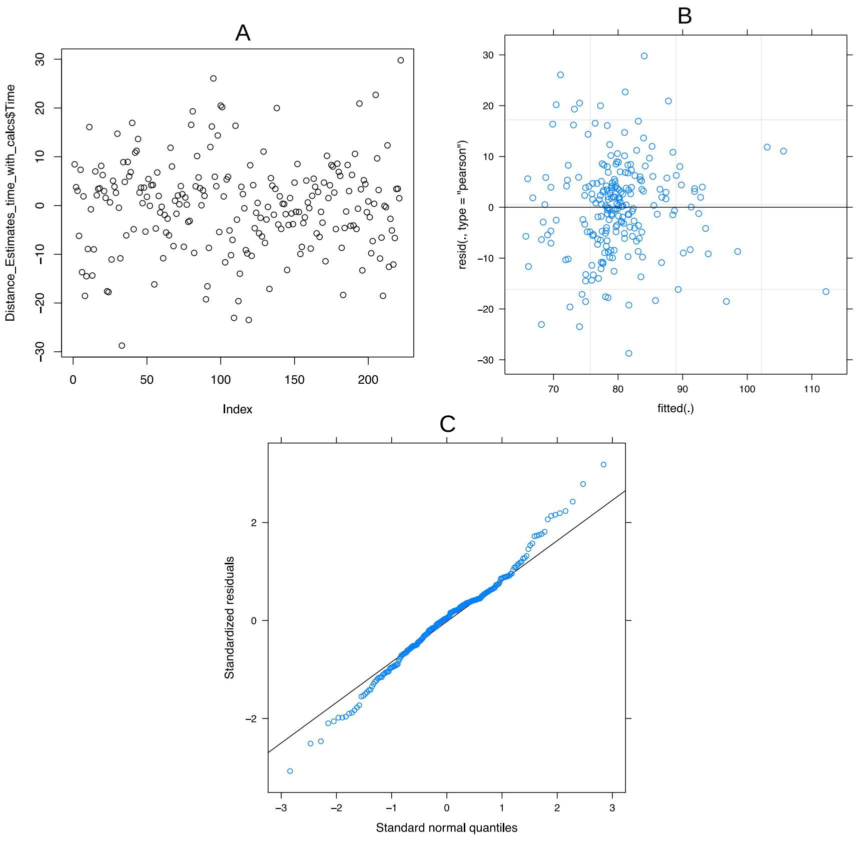


Figure 6: Testing mixed effects model assumptions (Absolute Turning Positon = Start Area Position + Fish Identity). (A): Plot of the model residuals against start area position. (B) A fitted vs residual plot revealing homogeneity of variance. (C)A QQ Plot of the standardised residuals, revealing a normal distribution.

##### 3.4 Linear mixed effects model: Distance Estimate = Speed + Fish Identity

The plot of residuals against the predictor (Swimming Speed) appears random (fig. 7, A), and we conclude the assumption has been met.

Variances were homogeneous, supported by the even distribution of residuals plotted against the fitted values (fig. 7, B).

A QQ-Plot of the standardised residuals indicates no major deviation from the straight line, and we conclude that they are therefore normally distributed (fig. 7, C).

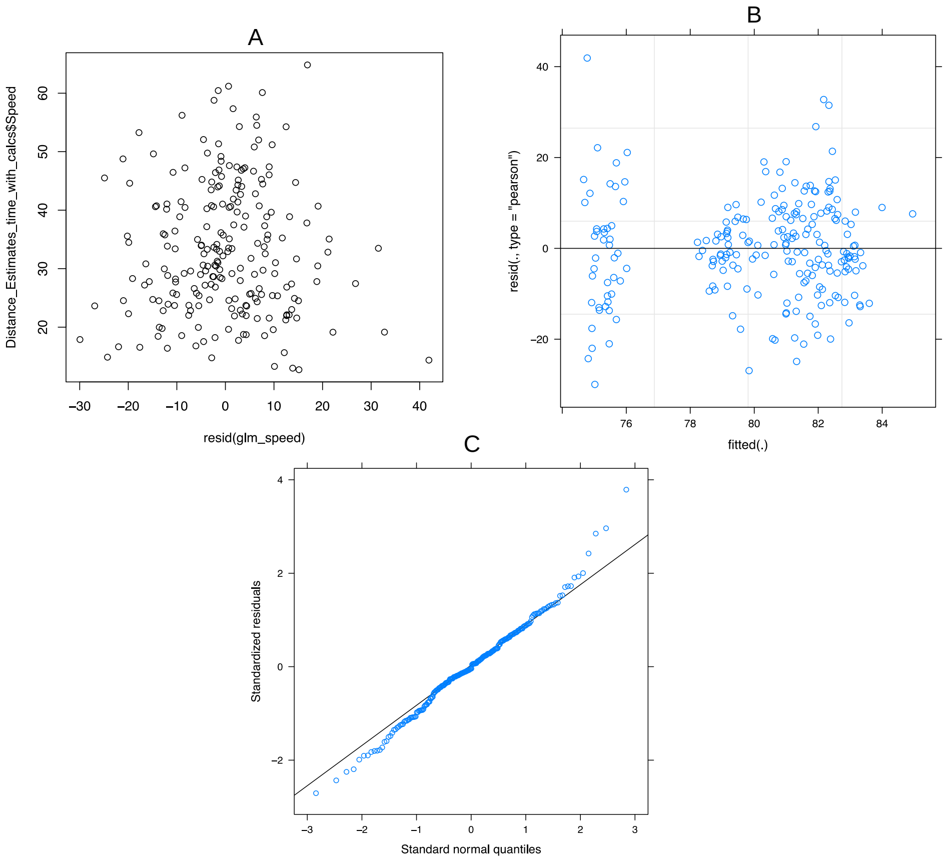


Figure 7: Testing mixed effects model assumptions (Absolute Turning Positon = Start Area Position + Fish Identity). (A): Plot of the model residuals against start area position. (B) A fitted vs residual plot revealing homogeneity of variance. (C)A QQ Plot of the standardised residuals, revealing a normal distribution.

#### 3.5 Linear mixed effects model: Distance Estimate = Tailbeat Number + Fish Identity

The plot of residuals against the predictor (Tailbeat number) once more appears random (fig. 8, A), and we conclude the assumption has been met.

Variances were homogeneous, supported by the even distribution of residuals plotted against the fitted values (fig. 8, B).

A QQ-Plot of the standardised residuals indicates no major deviation from the straight line, and we conclude that they are therefore normally distributed (fig. 8, C).

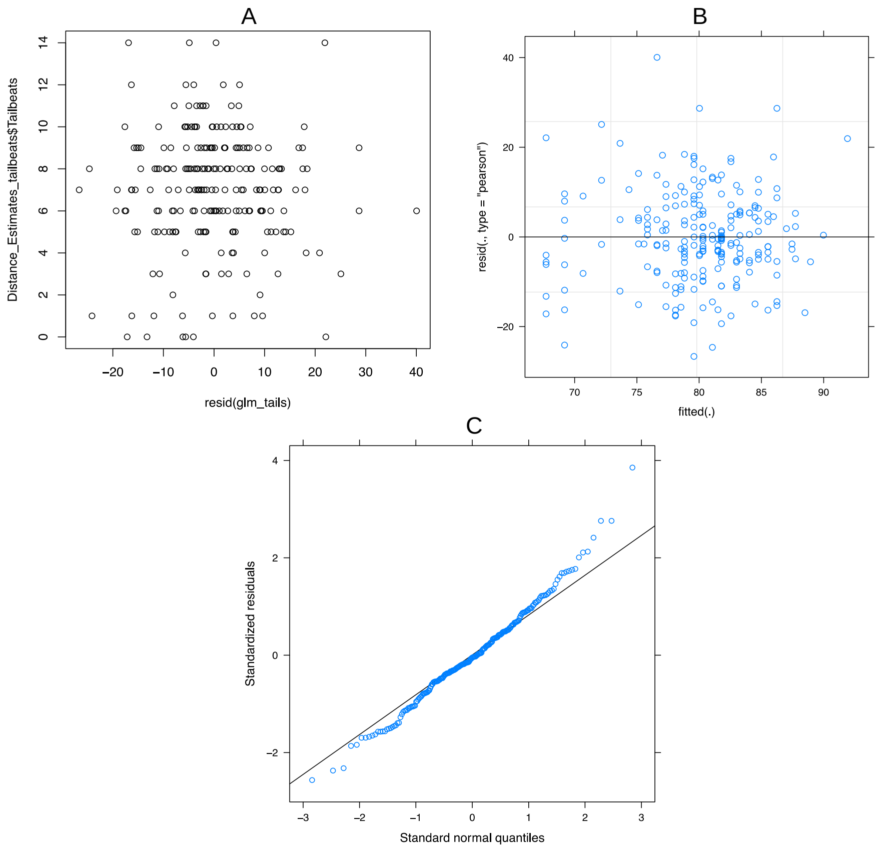


Figure 8: Testing mixed effects model assumptions (Absolute Turning Positon = Start Area Position + Fish Identity). (A): Plot of the model residuals against start area position. (B) A fitted vs residual plot revealing homogeneity of variance. (C)A QQ Plot of the standardised residuals, revealing a normal distribution.

### 4 Use of external cues for distance estimation

A linear mixed effects model was used used to assess whether fish were generalising across the three start area positions (+0, +10 and +20): Absolute Estimate Point = Start Area Position (Fixed Effect) + Fish (Random Effect). Start area position was a good predictor of the absolute estimate point within the tunnel across fish: $F_{210,2}$=58.8 (3sf), p<0.001, but some residual variation was explained by individual variation between fish (Likelihood Ratio Test with and without random effect = 7.63, p=0.00575 (3sf)). Figure 9 shows the absolute turning position for the three start area positions across all five fish. This confirms that our fish are not generalising across the start area positions, but are more likely to be using metric distance information.

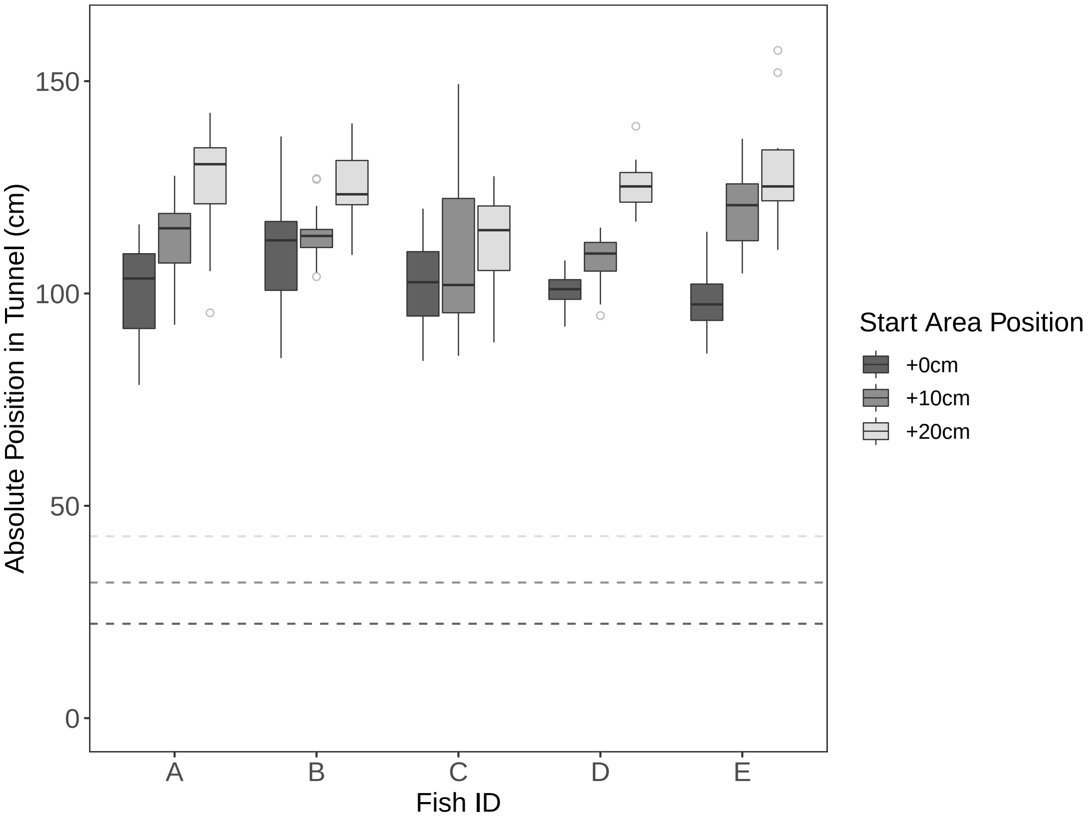


Figure 9: Absolute distance estimate position within the tunnel maze, split by start area position and fish identity. The start area moved between three positions: +0cm (dark grey), +10cm (mid-grey) and +20cm (light grey), all moving distally through the tunnel in 0.10m increments - the dashed lines indicate the position of the start area doorway from the back of the tunnel.
